# Error Correction Algorithms for Efficient Gene Expression Quantification in Single Cell Transcriptomics

**DOI:** 10.1101/2025.11.27.690682

**Authors:** Jens Zentgraf, Johanna Elena Schmitz, Andreas Keller, Sven Rahmann

## Abstract

Technological advances in single-cell RNA sequencing (scRNA-seq) allow us to sequence the transcriptomes of thousands of single cells in parallel, resulting in massive amounts of raw sequence data that must be processed efficiently to obtain a genes × cells expression matrix. In droplet-based scRNA-seq protocols, the sequenced mRNA molecules are tagged with a cell-specific barcode and a unique molecular identifier (UMI) within each cell. Both barcodes and UMIs may contain errors from production, amplification, or sequencing. Correcting and resolving such errors before further processing yields more reliable data and more accurate expression measurements.

We propose algorithmic advancements for barcode correction, read-to-gene mapping, and UMI resolution, which we combine into a new method called arcane for efficient gene expression quantification from scRNA-seq data. We additionally provide an implementation as a workflow-friendly command-line tool, also called arcane. This work builds on the recently published Fourway method to efficiently discover DNA *k*-mers with a Hamming distance of 1, speeding up barcode correction and UMI resolution and allowing for distinguishing *k*-mers into weakly and strongly unique ones during read-to-gene mapping. As a side result of separate interest, we show that for the mapping step, it suffices to store three genes per *k*-mer in order to cover almost all of the genes almost completely, thus avoiding arbitrarily large color sets in the colored De Bruijn graph index.

As a result, arcane is faster than existing methods while producing very similar results, as demonstrated in a comparison with CellRanger, Kallisto|bustools and Alevin-fry. arcane is available via GitLab (https://gitlab.com/rahmannlab/arcane).

## 1 Introduction

Over the past decade, single-cell and single-nucleus RNA sequencing (scRNA-seq, snRNA-seq) have become essential technologies for studying transcriptome heterogeneity across cells and for understanding rare cell types that were previously missed by bulk RNA sequencing [13,18,11]. For example, scRNA-seq has emerged as a key technology for cancer research since tumor heterogeneity poses a main challenge for understanding tumor development and successful treatment of cancer [5,24].

Droplet-based protocols, such as 10X Genomics (Chromium, chemistry v4), utilize two separate sequence tags to quantify the gene expression in thousands of single cells in parallel: the cell barcode and the unique molecular identifier (UMI) [15]. The sequencing adapters in each droplet have a unique cell barcode. Attached to each cell barcode is a UMI, such that each mRNA molecule inside a cell binds to a sequencing adapter with the same cell barcode but a different UMI. In this way, barcodes allow us to distinguish between different single cells, while UMIs are useful for deduplication of molecules of the same cell (after amplification) and unbiased counting.

The initial processing for scRNA-seq experiments consists of quantifying the gene abundance in each cell, which consists of three main steps. First, each sequenced fragment is assigned to its cell of origin, using its barcode. Second, the RNA/cDNA sequence of the fragment identifies the gene of origin. Third, fragments that share the same cell, gene, and UMI are clustered and collapsed into a single representative, so they are counted only once.

With error-free sequences, the first and third step would be straightforward. However, sequencing errors, production errors, and PCR amplification errors may lead to erroneous barcodes and UMIs, and hence to an inflation in distinct tags. Consequently, a critical step for accurate gene expression quantification is the implementation of robust barcode correction and UMI resolution methods, which are approached differently by existing tools and influence the quality of downstream analyses [19]. 10X Genomics provides a commercial solution, called CellRanger [31], to process their sequencing data. CellRanger performs splicing-aware alignment to the reference genome, which is computationally expensive. There is hence a high demand for faster methods. To fill this gap, alignment-free methods, such as Kallisto|bustools [16,22] and Alevin-fry [21,10] have been developed.

We introduce a new method, called arcane (**A**lignment-f**r**ee single **c**ell RN**A**-seq ge**n**e **e**xpression estimation) with algorithmic advancements that make arcane 2 to 3 times faster than the existing tools with very similar quantification results, but currently with higher memory requirements. The speed improvements mainly stem from the use of multi-way bucketed Cuckoo hash tables for fast gapped *k*-mer indexing [25,30] and, to a lesser extent, from a variant of a fast algorithm called Fourway [27,29] to efficiently discover pairs of sequences at Hamming distance 1 of each other, applied during barcode correction and UMI resolution. arcane comes with workflow-friendly command-line tools to quantify gene expression in single cell experiments. We now describe the existing approaches in more detail, focusing on barcode correction and UMI resolution. In Sec. 3, we describe the components of arcane in detail. We compare arcane with CellRanger, Kallisto|bustools and Alevin-fry in Sec. 4 and provide concluding remarks in Sec. 5.

## 2 Background and Related Work

We first explain the general steps of a scRNA-seq analysis workflow and how existing methods handle the computational challenges faced in each step.

Raw single cell RNA-seq data from droplet-based technologies consists of a stream of triples (*b, m, s*), where *b* is a DNA barcode identifying the cell, *m* is a unique molecular identifier (UMI), such that triples with the same (*b, m*) pair represent copies of the same molecule from the same cell, and *s* is the RNA sequence that allows us to identify the gene of origin. In practice, this information may be distributed across different files. For 10x Genomics data, read 1 (R1) contains both barcode and UMI, and read 2 (R2) contains the cDNA sequence of the (m)RNA molecule. Typical lengths are |*b*| = 16, |*m*| = 12 and |*s*| ≈ 100 bp. We assume that a positive list *B* of known valid barcodes exists.

To generate a genes × cells expression matrix, the three main steps are barcode correction, gene (or transcript) identification, and UMI resolution. For both barcode correction and UMI resolution, we assume that the sequences have reasonable error rates and that errors are mostly substitutions. Then, of all the barcodes from a cell, the correct barcodes will be the most frequent ones^1^. Among the erroneous variants, the most frequent ones are those with Hamming distance 1 from a correct barcode. The same assumption holds for UMIs.

### 2.1 Barcode correction

The first step for creating a gene expression count matrix for 10x Genomics sequencing data is to sort the sequenced fragments by cell of origin, which means grouping reads with the same barcode. As the barcode may contain errors (from production, amplification, or sequencing), one needs a method to deal with invalid observed barcodes (those that are not on the positive list *B* of valid barcodes). Since valid barcodes have a high abundance, the simplest solution is to only consider barcodes from *B* that appear frequently in the data (above a cutoff). However, this approach may discard a substantial fraction of reads that still contain useful information. Thus, many approaches correct invalid barcodes into a barcode from *B*, e.g., if they have a (Hamming or edit) distance of 1.

Next, among the remaining distinct barcodes, those of low abundance are declared as noise and ignored. Such low-abundance barcodes may result from droplets with no cell or a mixture of cells. Instead of using a fixed abundance cutoff, Alevin-fry, Kallisto|bustools and CellRanger set the threshold by finding a “knee” in the cumulative distribution function of the total associated read counts (or UMI counts) of each barcode (see Sec. 3.3).

### 2.2 Gene identification

In the next step, the sequence reads are mapped to a gene or transcript. For this, CellRanger performs a splicing-aware alignment of each read to the genome using the STAR aligner [6]. The gene is then identified based on the position it was mapped to, using transcript annotation information from a GTF file. Due to the high computation time of alignments, Alevin-fry [10] and Kallisto|bustools [16] perform a pseudo-alignment using a colored de Bruijn graph [12]. In a de Bruijn graph, every node represents a *k*-mer and there exists a directed edge between two nodes *q, q*^′^, if the last *k* − 1 positions of *q* and first *k* − 1 positions of *q*^′^ are equal [4]. In a colored de Bruijn graph, each node is additionally annotated with a set of colors. For gene expression quantification, a “color” means a gene (or transcript) that a *k*-mer occurs in [2]. Thus, we can map a read to a gene by finding a path in the de Bruijn graph where the color sets of all nodes contain the same gene. In practice, colored de Bruijn graphs are implemented using hash tables that support fast *k*-mer queries instead of storing the graph structure explicitly [2,1,17]. Both Alevin-fry [10] and Kallisto|bustools [16] combine pseudoalignment with a probabilistic approach to quantify gene abundances, so they are able to take into account information from reads that map equally well to different genes or transcripts.

### 2.3 UMI resolution (De-duplication)

For accurate gene expression quantification, we need to remove UMI PCR duplicates to ensure that each sequenced RNA molecule within the same cell is counted only once after amplification. The simplest solution, as done by Kallisto|bustools, is to collapse all reads with the same UMI, barcode, and gene. However, this may artificially inflate gene counts, since for each UMI with a (sequencing) error, the corresponding gene is counted multiple times within the cell. We hence have to solve the problem of distinguishing correct UMIs from spurious UMIs that arise due to errors.

To circumvent this problem, CellRanger only considers UMIs that have no undetermined bases (`N`s), that are not homopolymers, and that have a sequencing quality ≥ 10 for every base. In addition, all UMIs with an edit distance of 1 are collapsed into a single UMI. This can lead to a transitivity problem, which has to be considered during UMI resolution. For example, if an error occurs during PCR and another error is introduced during sequencing, we have multiple pairs of edit distance 1 neighbors that should be collapsed. Previous work has shown that this approach tends to underestimate the total number of distinct UMIs [20], and therefore more advanced UMI resolution algorithms have been developed.

UMI-tools [20] proposes several graph-based approaches. The approach with the highest accuracy builds a graph (separately for each cell) from all UMIs mapping to the same position in the reference genome. Each node corresponds to a distinct UMI and there is a directed edge between two nodes *u* and *v* if their UMIs have an edit distance *< T* and if *n*_*u*_ ≥ 2*n*_*v*_ − 1, where *n*_*u*_ and *n*_*v*_ are the UMI counts of nodes *u* and *v*, respectively. The UMIs of a weakly connected directional network are then collapsed.

Alevin-fry supports several different UMI resolution modes that are either similar to Cellranger or build a parsimony graph to find the minimal number of molecules that can explain the graph (considering the UMI sequence, UMI count, and transcript mapping).

## 3 Accurate and Efficient Gene Expression Quantification (arcane)

After giving basic definitions and stating fundamental algorithms, in particular the Fourway algorithm for finding pairs of sequences at Hamming distance 1 (Sec. 3.1), we describe arcane’s index creation step for assigning genes to reads (Sec. 3.2 and Fig. 2). Then, we discuss the three main steps of arcane’s gene expression quantification method (overview in Fig. 3): barcode correction (Sec. 3.3), assignment of reads to genes (Sec. 3.4), and UMI resolution (Sec. 3.5). Implementation details, such as parallelization and use of shared memory, are described in Appendix F.

### 3.1 Fundamental Definitions and Algorithms

#### DNA k-mers and encoding

The raw sequencing reads are nucleotide sequences over the DNA alphabet *Σ* = {`A, C, G, T`} . Since the alphabet size is |*Σ*| = 4, we encode a single nucleotide by log_2_ |*Σ*| = 2 bits using the encoding `A` ↦ (0)_4_ = (00)_2_, `C` ↦ (1)_4_ = (01)_2_, `T` ↦ (2)_4_ = (10)_2_ and `G` ↦ (3)_4_ = (11)_2_. A *k-mer* is a string of length *k* over *Σ*, i.e., an element of *Σ*^*k*^. The encoding *enc*(*x*) for a *k*-mer *x* reads *x* as a base-4 number using the above encoding, concatenating from left to right. For *k* ≤ 32, *enc*(*x*) can be stored in a single 64-bit integer. Given two integers 0 *< k* ≤ *w*, a gapped *k*-mer is a string of length *k* that considers *k* positions in a window of length *w*, specified by a *mask*. A (*k, w*)-mask is a string of length *w* over the alphabet {`#,_`} that contains exactly *k* times the character `#` and *w* − *k* times the character `_`. The *k* positions with the `#` character are called *significant positions* of the mask. Equivalently, the significant positions may be specified by the *k*-subset (or sorted tuple) of significant positions. It is customary to require that the mask starts and ends with `#`; otherwise, non-significant positions at either end may be removed [14,3,28].

#### Rank queries

Given a (cache-line aligned) bit array *A*[0 … (*n* − 1)] with *n* bits, we want to answer rank queries, i.e., count the number of ones (or zeros) up to a given index, fast and without requiring much extra space. For 0 ≤ *i* ≤ *n*, the 1-rank of position *i* in *A* is defined as

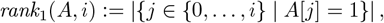

and accordingly *rank*_0_(*A, i*) := |{*j* ∈ {0, …, *i*} | *A*[*j*] = 0 }|= (*i* + 1) − *rank*_1_(*A, i*) for zeros. We use a fast, simple single-level rank data structure [8]: Assuming *n* ≤ 2^32^, we use an array of pre-computed 32-bit ranks *R*[*i*] := *rank*_1_(*A*, 512 *i* − 1) at every 512-th bit; this needs 32*n/*512 = *n/*16 bits (6.25%) of additional space. To answer general rank queries *rank*_1_(*A, i*), we look up *R*[⌊*i/*512⌋] and count the (at most 511) remaining bits that are within a single cache line using elementary `popcount` CPU instructions on native 64-bit integers, in combination with bit-masks.

#### The Fourway algorithm for Hamming-distance-1 neighbor pairs [29]

During several of arcane’s steps, we need to identify all pairs of sequences from a given large set *X* of *k*-mers (with *k* ≤ 32) that have a Hamming distance of 1. This also applies to gapped *k*-mers once they have been reduced to a sequence of length *k*. Naively comparing all pairs is typically impractical, with *X* containing several millions or billions of sequences. One approach is to store the set *X* as a hash table for constant-time membership tests, iterate over *x* ∈ *X*, create the 3*k* neighbors of *x* at Hamming distance 1 and test each of them for presence in *X*. If |*X*| = *n*, the naive pairwise testing takes *O*(*n*^2^) time, while the neighborhood generation takes only *O*(*nk*) time, which is a considerable improvement. Recently, we developed a method called the Fourway algorithm [29] that is even faster in practice, even though the asymptotic time is also *O*(*nk*). We now give a summary of the Fourway algorithm, taken from [27,29].

We assume that the *k*-mers are available in a lexicographically sorted array *A* of length *n*. The algorithm is similar to a 4-way merge and proceeds recursively. It is called as Fourway(*A, d, q*) with a depth parameter *d* ∈ {1, …, *k*} and a list of start pointers *q* = (*q*_*c*_) with *c* ∈ {`A, C, G, T, $`}. The invocation Fourway(*A, d, q*) searches for Hamming-distance-1 pairs that disagree exactly at the *d*-th nucleotide among the *k*-mers in the interval *I* := [*q*_A_, *q*_$_ [of *A*, in which the first (*d* − 1) nucleotides of the contained *k*-mers are equal, *q*_*c*_ (for each nucleotide *c*) points to the start of the sub-interval of *I* where the *d*-th nucleotide is *c* (Fig. 1) and the sentinel pointer *q*_$_ points beyond the end of *I*. In the initial call, the depth is *d* = 1 (no common prefix, the first nucleotide may differ), *q*_A_ = 0 points to the first (smallest) element in *A*, and *q*_$_ = *n* points past the end of A. The initial values of the other pointers *q*_*c*_, *c* ∈ {`C, G, T`} are determined by linearly scanning over *A* once.

**Fig. 1:**
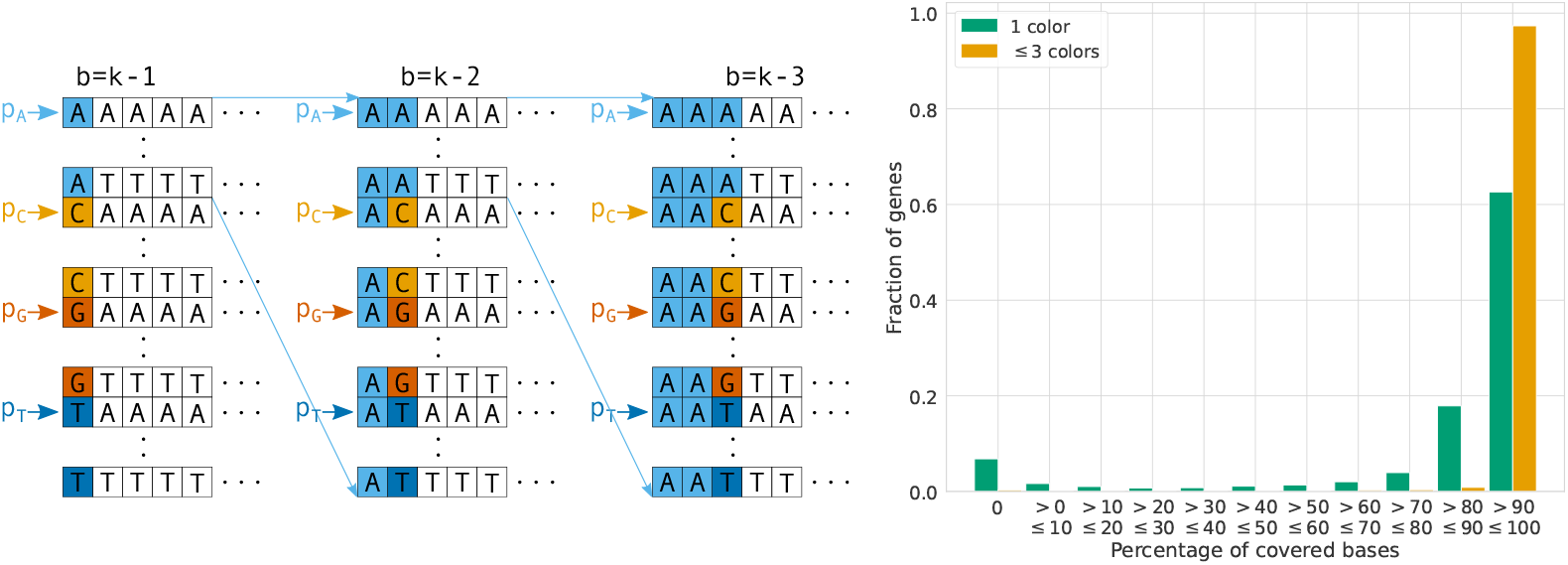
**Left:** Recursive 4-way comparison at depths *d* = 1, 2, 3, from left to right. At depth *d*, the first *d* − 1 characters of all *k*-mers within the considered interval are equal, and the *d*-th character is compared. Let *b* := *k* − *d*. The (at most) four *b*-mer suffixes of the *k*-mers pointed to by pointers *p*_*c*_, *c* ∈ *Σ* are compared, and the lexicographically smallest *b*-mers are identified among the four elements. The *k*-mers with equal smallest *b*-mers are marked as weak if there are at least two equal minimal *b*-mers. After the 4-way comparison of an interval is completed, it is repeated recursively with increased depth *d* + 1 on its 4 sub-intervals (only the first recursive call for the `A`-subinterval is shown). **Right:** Color information for the human transcriptome (using CellRanger reference and GTF file; containing transcripts and intronic regions). The plot shows the fraction of genes (*y*-axis) that have a given percentage (*x*-axis) covered by either unique gapped *k*-mers (green) or by gapped *k*-mers appearing in up to three genes (orange).

**Fig. 2:**
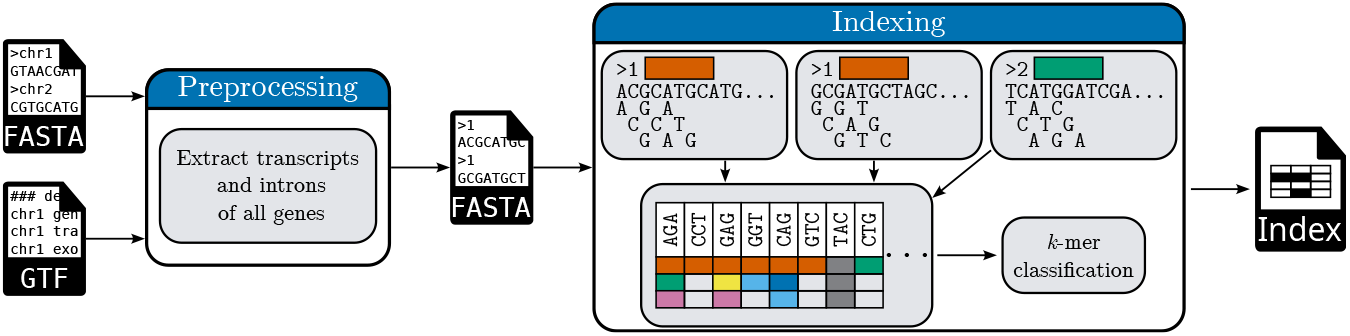
Creation of a gapped *k*-mer index. First, the reference genome (FASTA format) and an associated annotation file (GTF format) are processed to create a new FASTA reference file that contains all transcripts and introns for all annotated genes. The header of each sequence contains the numerical gene identifier; several sequences (transcripts, introns) in the file may belong to the same gene. In the indexing step, all sequences are split into their overlapping gapped *k*-mers, and a hash table is constructed that maps each *k*-mer to the set of identifiers of the genes in which the *k*-mer occurs. After inserting all *k*-mers, we classify them into strongly unique *k*-mers, weakly unique *k*-mers and non-unique *k*-mers (see Section 3.2).

**Fig. 3:**
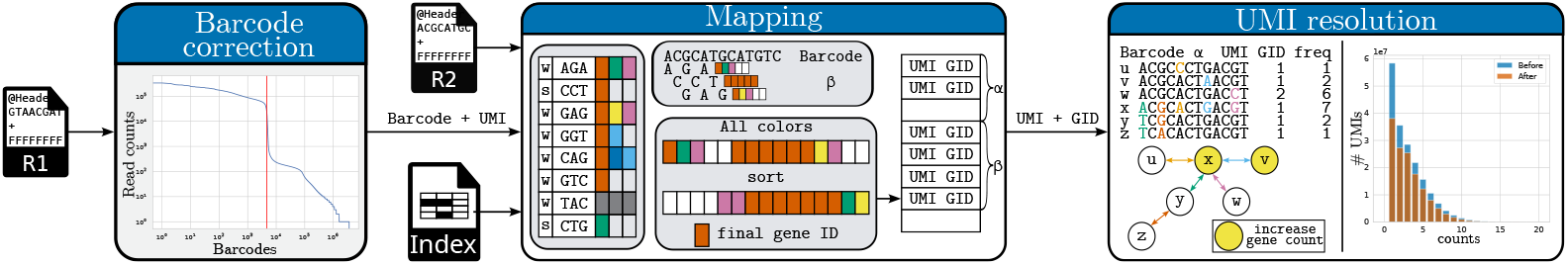
Overview of arcane’s gene expression quantification steps. arcane first performs barcode correction and removal of empty droplets. Next, all reads with a valid (and possibly corrected) barcode are mapped to the transcriptome using a pre-built gapped *k*-mer index that maps gapped *k*-mers to genes. The third step is UMI resolution; arcane quantifies the gene counts in each cell after collapsing UMIs that likely belong to the same RNA molecule.

First, we create working copies *p* = (*p*_*c*_) of *q* = (*q*_*c*_) for *c* ∈ *Σ*. While *q* will stay unchanged, the *p*_*c*_ increase towards larger elements as the algorithm proceeds. Initially, all the *p*_*c*_ pointers are *active*. When *p*_*c*_ moves beyond the end of its sub-interval (i.e., it reaches a *q*_*c*_*′* for a character *c*^′^ *> c*, where the sentinel is larger than any character in *Σ*), the pointer becomes *inactive*. If only a single pointer or no pointer is active, we are done and proceed to recursive calls at larger depth; see below.

In each step, the algorithm examines the *k*-mers at the locations pointed to by the active pointers *p*_*c*_. We call these the *active k-mers*. They jointly have the following properties: (1) their first *d* − 1 characters are equal (true for all *k*-mers in interval *I*), (2) their *d*-th characters are distinct, (3) their suffixes of length *b* = *k* − *d* are arbitrary, but examined in increasing order, and (4) they are the smallest *k*-mers in *I* that have not yet been (but still may be) identified as weak based on a single difference at their *d*-th character. We look at the *b*-suffixes of the active *k*-mers and find the smallest one(s). Let *C*^∗^ be the character set such that the *k*-mers at *p*_*c*_, *c* ∈ *C*^∗^, have the minimal *b*-suffixes among the active *k*-mers. If |*C*^∗^| ≥ 2, we have identified a group of *k*-mers of size |*C*^∗^| that differ only at their *d*-th position; hence all of them are marked as weak. (If |*C*^∗^| = 1, nothing happens.) Then, all *p*_*c*_ for *c* ∈ *C*^∗^ are incremented. These steps are repeated until a single (or no) active *p*_*c*_ remains.

After processing interval *I*, if *d < k*, the sub-intervals are processed recursively, so there are |*Σ*| = 4 recursive calls, each with increased *d* ← *d* + 1 (and reduced *b* = *k* − *d*). The initial pointers *q* for each subinterval are obtained by a linear scan through the sub-interval. The recursive call is not performed if the length of the subinterval is at most 1, as there is nothing to compare then.

The original publication [29] describes several optimizations of this basic idea, which we also use, and additionally considers the case where a *k*-mer is treated as equivalent to its reverse complement, which we do not need because the sequences in arcane are strand-specific.

### 3.2 Creation of arcane’s gapped *k*-mer index

For gene identification from the RNA reads, arcane requires a species-specific gapped *k*-mer index that stores for each *k*-mer the genes in which it occurs, also referred to as colors^2^. For each species, the index creation needs to be executed only once.

#### Preprocessing

Before arcane builds the index, it takes a reference genome in FASTA format and a GTF file with gene annotations to produce a new FASTA file. Currently, arcane supports filtered GTF files produced by CellRanger that contain only the relevant gene entries for gene expression quantification. The new FASTA file contains all known transcripts, introns, and exon-intron junctions on the correct strand for all protein-coding genes and additional biotypes, such as lncRNA. For the exon-intron junctions, we add an overlap of (*w* − 1) bp to the intron size at the start and end of the intron in order to guarantee that we do not lose any gapped *k*-mers (of width *w*) that appear in reads of unspliced transcripts. Every gene is assigned a unique 16-bit integer ID, which is stored in the header lines of the FASTA file with all sequences derived from that gene. A separately stored table of Ensembl gene IDs (ENSG…) and gene names maps the numerical gene ID to an interpretable string.

#### Index creation

Given the preprocessed FASTA file with gene ID headers, we insert all gapped *k*-mers into a 3-way bucketed Cuckoo hash table [25,26,30], which stores the set of genes each gapped *k*-mer occurs in. Since 10X Genomics data is strand-specific, arcane currently only stores the gapped *k*-mers on the sense strand and does not support antisense mapping.

Since we store the colors directly with the gapped *k*-mer without another memory indirection, we have to limit the number of colors. Note that other approaches solve this problem by storing the color sets in an auxiliary data structure, which usually causes one or more additional cache misses. If we only consider gapped *k*-mers with a single color, only 62.65% of all genes have *>* 90% of their positions covered by at least one such *k*-mer in the human reference. If we increase the limit of the number of colors per *k*-mer from 1 to 3, then 97.3% of the genes are covered at *>* 90% of their positions (Fig. 1, right). Therefore, we store up to three colors for each gapped *k*-mer. If a gapped *k*-mer occurs in more than three genes, we store a special value Multi, indicating that the number of genes is too large to be stored.

If for a gapped *k*-mer, a single nucleotide variation could change one of the assigned genes, the gene mapping is less robust (see also Section 3.4). Therefore, we store an additional bit, called the “weak bit”, to indicate whether a Hamming-distance-1 neighbor of the *k*-mer exists with a color set that is not a superset of the *k*-mer’s color set. All pairs of gapped *k*-mers at Hamming distance 1 are discovered using the Fourway algorithm [29], as described in Sec. 3.1. A gapped *k*-mer is called *non-unique* in the stored table if it has more than one color; otherwise, it is *unique*. We further distinguish between two types of unique gapped *k*-mers: Unique *k*-mers with their weak bit set are less reliable, and a single nucleotide change can change their color set, so we call them *weakly unique*. Otherwise, they are *strongly unique*.

Since Multi values contain no helpful information, we reduce the index size by only storing the gapped *k*-mers with ≤ 3 colors. For this, we count how many gapped *k*-mers in the index satisfy this property, copy them to a new precisely sized smaller hash table, and forget the original table.

### 3.3 arcane’s barcode correction algorithm

Given a positive list *B* of valid barcodes, arcane first identifies erroneous variants of valid barcodes and attempts to correct them, where possible, to the best-fitting valid barcode.

*Data structure*. We represent each barcode *b* that is observed at least once in the dataset by a numerical index *j*_*b*_, preserving the lexicographic order of the barcodes, i.e., *j*_*b*_ *< j*_*b*_*′* if and only if *b < b*^′^. For this, we use a bit array *A* of 4^*ℓ*^ bits, where *ℓ* is the barcode length (*ℓ* = 16 in our data, so *A* uses 4 Gbits or 0.5 GB)^3^. Initially, all bits are set to value 0. If a barcode *b* appears anywhere in the dataset, we set bit *enc*(*b*) to 1.

More concretely, we stream all R1 reads of the dataset, extract the barcode *b*, compute *enc*(*b*) and set the bit. We build the rank data structure described in Sec. 3.1 and define *j*_*b*_ := *rank*_1_(*A, enc*(*b*)) − 1 . Let *c* be the number of 1-bits in *A*, i.e., the number of distinct barcodes in the sample. Then, the definition of *j*_*b*_ bijectively maps the indices of these *c* bits in *A* to {0, …, *c* − 1} in an order-preserving way. We store information about how to correct barcodes in an array *C* of size *c*, such that information about barcode *b* is stored at *C*[*j*_*b*_]. This information is a 4-tuple (*enc*(*b*), *valid*(*b*), *count*(*b*), *correct*(*b*)), where

1. *enc*(*b*) is the base-4 encoding of *b*, representing *b* itself. In principle, *enc*(*b*), does not need to be stored, as it can be obtained by a select query on *A* (given *j*, find the index *e* = *enc*(*b*) of the *j*-th 1-bit in *A*; inverse to rank), but prioritizing speed over memory, we store *enc*(*b*) explicitly at *C*[*j*_*b*_].
2. *valid*(*b*) is a single bit that indicates whether *b* is a valid barcode from the positive list *B*;
3. *count*(*b*) is the number of reads with barcode *b*;
4. *correct*(*b*) is the index *j*_*b*_*′* of the barcode *b*^′^ to which *b* should be corrected. Initially *correct*(*b*) = Unknown, a special value, if *b* is not valid, and *correct*(*b*) = *j*_*b*_ if *b* is valid.

#### Correction algorithm

We use the Fourway algorithm [29], as described in Sec. 3.1 to discover all pairs of present barcodes that have a Hamming distance of 1. Whenever Fourway discovers a pair {*b, b*^′^} of barcodes with Hamming distance 1, we do the following.

If none or both of *b, b*^′^ are on the positive list *B* of barcodes, nothing happens, as none can be corrected into the other. If exactly one of them, say *b*, is on the positive list (*valid*(*b*) = 1), then *b*^′^ may eventually be corrected into *b*. If *correct*(*b*^′^) is still Unknown, we set *correct*(*b*^′^) ← *j*_*b*_ to save *b* as a valid candidate for *b*^′^. However, if *correct*(*b*^′^) already has a different value *b*^0^ ≠ *b*, then there are at least two Hamming-distance-1 neighbors of *b*^′^ in *B*, and we cannot make a reliable decision. In that case, we set *correct*(*b*^′^) ← Ambiguous and never change it again.

At the end of the Fourway algorithm, we execute all feasible corrections: For each *b*^′^ with *correct*(*b*^′^) =*b* ∉ {*b*^′^, Unknown, Ambiguous}, we add *count*(*b*^′^) to *count*(*b*) and set *count*(*b*^′^) ← 0.

#### Removal of infrequent barcodes

After the barcodes have been corrected and their counts updated, arcane supports three options to remove barcodes that occur infrequently and likely do not represent actual cells.

1. If the number *c* of sequenced cells is provided by the user, the *c* most frequent barcodes from the positive list are kept and the remaining ones are discarded.
2. We detect the “knee” in the complementary cumulative distribution function (ccdf) of the barcode counts using the implementation of UMI-tools [20]; see Appendix A and its Fig. A1 for details. In distance mode (default), the knee is defined as the point in the cumulative distribution where the distance between the cumulative distribution curve and a straight line between the first and last points on the cumulative distribution curve is maximal. All barcodes above the knee are assumed to represent cells.
3. Alternatively, all barcodes on the positive list may be kept (even if they are infrequent), which may be of interest if downstream processing includes batch correction tools that require empty droplets to estimate the background noise [7].

#### Output

The final result is an array with information about the kept valid barcodes and the correctable barcodes. Each element stores a triple (*enc*(*b*), *valid*(*b*), *info*(*b*)), where *enc*(*b*) and *valid*(*b*) is as above, and *info*(*b*) = *count*(*b*) if *b* is a valid barcode, but *info*(*b*) = *correct*(*b*) if *b* is a correctable barcode. For each barcode *b*, the triple fits into a single 64-bit integer (32 bits for the 16 bp barcode, 1 bit for *valid*(*b*), and 31 bits for *info*(b)). The discarded barcodes do not appear.

### 3.4 Mapping reads to genes

The input for the mapping step consists of the barcode correction information and of a stream of triples (*b, m, s*), where *b* and *m* are the barcode and UMI associated with the RNA sequence *s*, respectively. The goal is to find the gene of origin of sequence *s*, represented by the gene identifier *g* = *gid*(*s*). We directly discard all triples with an invalid barcode and where barcode or UMI contain an undetermined base (`N`).

For each remaining triple, we correct the barcode if necessary, and then query all gapped *k*-mers of the read *s* in the index, retrieving the associated numeric gene identifiers for each *k*-mer, or the information that the *k*-mer is not present. We collect weighted gene identifier information of each gapped *k*-mer of the read in a list, as we iterate over the read. The (two or three) gene IDs of a non-unique *k*-mer are counted with weight 1, the (single) gene ID of a weakly unique *k*-mer with weight 3, and the (single) gene ID of a strongly unique *k*-mer with weight 5. Given the final list, we sort it and sum the weights of each gene identifier and search for the most frequent one (say, *g* with count *q*) and for the second most frequent one (say, *g*^′^ ≠ *g* with count *q*^′^ *< q*). If such a *g* or *g*^′^ does not exist, we use *q* = 0 or *q*^′^ = 0, respectively. If *q* ≥ *q*^′^ + 3, we assign the read to the gene with identifier *g*, i.e., we set *gid*(*s*) := *g*. Otherwise, we declare the read as not (reliably) mappable, *gid*(*s*) := Unmapped.

To store the mapping results, we use a large array, divided into blocks, with one block per corrected barcode (cell). The size of each block is known after the barcode correction step, so the array can be allocated with the correct size. Each element of the block for barcode *b* stores information about one read with corrected barcode *b*. This information consists of the (so far uncorrected) UMI *m* and of the gene identifier *gid*(*s*), which may be Unmapped. Thus, for a triple (*b, m, s*), we identify the first free position in the block for barcode *b* in the output array, and store (*m, gid*(*s*)) there.

### 3.5 UMI resolution algorithm

The input for UMI resolution is one block of the output of the read-to-gene mapping step (Sec. 3.4), i.e., a list of (UMI, gene id) pairs (*m, g*). UMI resolution thus proceeds separately for each barcode and potentially in parallel over barcodes (cells). The goal is to count each gene only once for each (correct) UMI, collapsing multiple occurrences of the same corrected UMI for the same gene. The final output is a genes × cells expression count matrix *X* = (*X*_*g,b*_) (in a sparse matrix format).

For a fixed barcode *b*, the first step is to count and collapse the list of (*m, g*) pairs into a mapping of type `UMI → List[Pair[GeneID, Count]]`, such that each UMI is mapped to a list of (gene, count) pairs that indicate which gene(s) are seen how many times with UMI *m* in the input list. The second step is to run Fourway on the sorted UMIs to build (conceptually) a graph, where the nodes are the UMIs and an edge exists between UMIs *m m*^′^ with Hamming distance 1. We then determine the connected components (UMI clusters) of this graph. In the implementation, we do not create the graph explicitly, but combine Fourway with a Union-Find data structure [23] to maintain a partition of the set of UMIs. arcane introduces a new UMI resolution strategy, called network mode, to decide which gene(s) to count in each component. For a comparison between UMI counts before and after UMI resolution using network mode and previously suggested strategies, see Appendix B.

#### Network mode

To decide if two UMIs within the same connected component should be collapsed into a single UMI or not, we first estimate the expected number of reads for a single UMI after PCR duplication. We assume that the number of reads per UMI follows a zero-inflated Poisson distribution. The expected UMI count is thus given by the parameter *λ* of a Poisson distribution. To avoid a biased estimate due to high singletons (UMIs with a read count of 1, partially due to sequencing errors) and some outliers with unusually high counts, we consider the relation of *λ* to the ratio *P* [*Z* = 3]*/P* [*Z* = 2] = (*λ*^3^e^−λ^*/*6)*/* (*λ*^2^e^−λ^*/*2) = *λ/*3 for a Poisson(*λ*) random variable *Z*.

For a sample of *N* UMIs, we approximate *P* [*X* = *k*] by *f*_*k*_*/N*, where *f*_*k*_, is the number of UMIs that are seen *k* times in the sample. We thus estimate 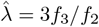, separately for each barcode (cell). We apply the following UMI resolution rules within a connected component of the graph discussed above.

1. For all UMI-gene combinations with a count 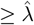, we count the gene once. Therefore, a connected component can count the same gene multiple times to mitigate over-collapsing of UMIs.
2. If, for a gene, no UMI is above the threshold, we count the gene once if the sum over all UMIs associated with this gene is 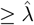.
3. If no gene was incremented based on the above two rules, but there is only a single gene in the connected component, we count that gene once. An isolated UMI-gene combination with a small count is most likely a true UMI due to the low coverage in scRNA-seq data.

## 4 Evaluation Results

We compare the gene expression counts and resource usage of arcane with those of CellRanger (version 9.0.1), Kallisto|bustools (version 0.30.0) and Alevin-fry (version 0.11.2) on three public 10x Genomics Chromium human datasets and on one mouse dataset: The `5K_Human_PMBC_v3` and `10K_Human_PMBC_v3` datasets contain approx. 5 000 resp. 10 000 human peripheral blood mononuclear cells (PBMCs), processed with v3 chemistry. The `10K_Human_Melanoma_v4` dataset contains approx. 10 000 disseminated tumor cells from a melanoma cancer taken from a human donor, processed with v4 chemistry. The mouse dataset (`10K_Mouse_Brain_v4`) is the only single nucleus RNA-seq dataset; it contains approx. 10 000 mouse brain cells and was processed with v4 chemistry. URLs for the datasets are provided in Appendix E.

We ran all tools using the default parameters for their respective versions, if not explicitly stated otherwise. All tools are configured to output merged gene counts (spliced and unspliced). For Alevin-fry, the unspliced, spliced, and ambiguous counts (USA mode) are summed up for each gene. We first created *k*-mer indices for arcane, Kallisto|bustools and Alevin-fry. For arcane, we used the (31, 43) mask `####_#_##_###_#_###_###_###_#_###_##_#_####` to build the human and mouse index. It guarantees at least 4 hits (and 51 covered positions) in a read of length 100 with at most 3 substitution differences compared to the reference [28]. As Kallisto|bustools supports only contiguous *k*-mers, we used the default of *k* = 31, and Alevin-fry was run with default *k*-mer length of 31 and minimizers of default length 19. We provided the same input files to all tools, namely the latest genome reference sequences and the GTF annotation file generated with CellRanger. For Kallisto|bustools, the index is built with `--workflow=nac` to be able to map both nascent and mature mRNA molecules. Since Kallisto|bustools has no built-in step to remove rare barcodes, we calculate the “knee” as a post-processing step using UMI-tools [20]. We ran Alevin-fry using the SIMPLEAF wrapper (version 0.19.5) [9] using the `--knee` option to filter barcodes and setting the UMI resolution mode to either `cr-like` or `parsimony-em`. For arcane, we used the `network` mode described in Sec. 3.5.

All benchmarks were run on an Ubuntu (24.04 LTS) server with two AMD EPYC 9534 64-Core Processors, 1.5TB of 4800-MHz DDR5 memory and a KIOXIA CD8P SSD. Running time and maximum memory usage were measured using `/usr/bin/time -v`.

### 4.1 arcane is fastest with larger memory requirements

We compare the computational resource requirements of arcane with those of the other tools. Fig. 4 shows the wall clock running times and memory usage of arcane, Cellranger, Kallisto|bustools and Alevin-fry (using both `cr-like` and `parsimony-em` UMI resolution modes).

**Fig. 4:**
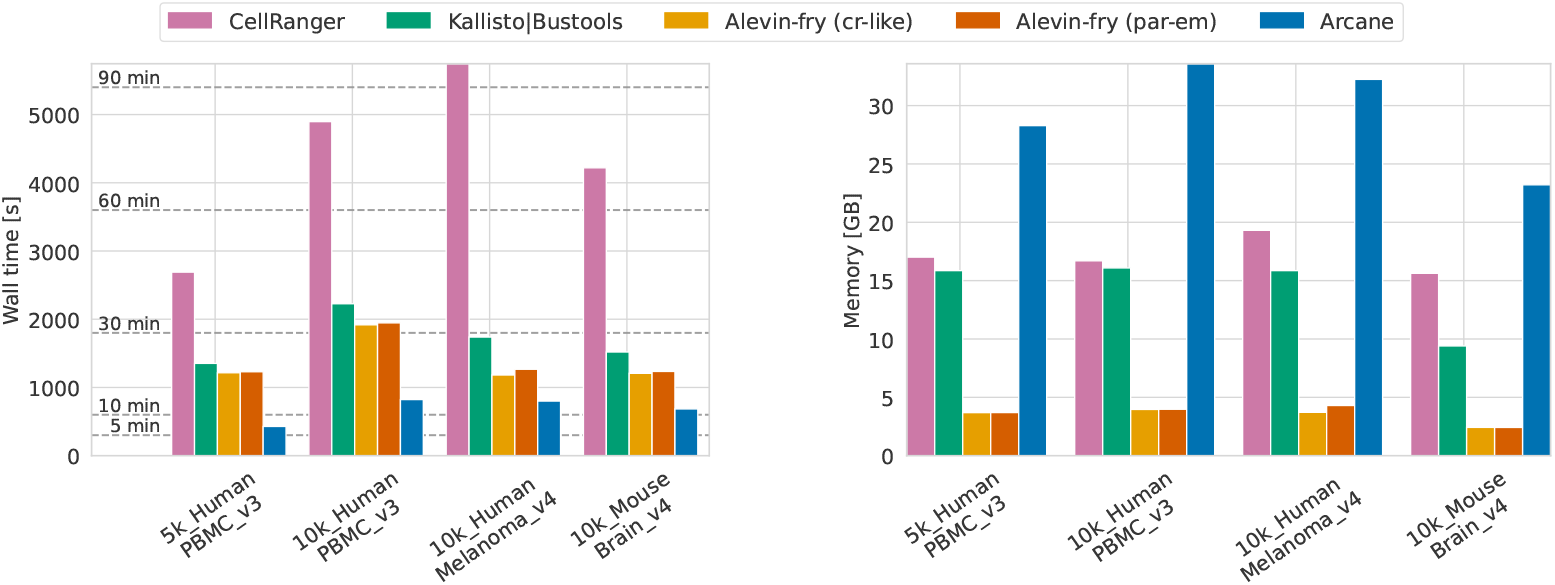
Comparison of wall time and memory usage across tools for all four datasets using 16 threads.

On all four datasets, arcane is fastest, taking under 13 min on each dataset. Alevin-fry and Kallisto|bustools have similar running times (e.g., 32 min and 37 min on `10K_Human_PBMC_v3`, respectively). In contrast to the other approaches, the mapping step of CellRanger is alignment-based, and hence CellRanger is by far the slowest tool on all datasets, e.g. 96 min on `10K_Human_Melanoma_v4` vs. 29 min for Kallisto|bustools, 20 min for Alevin-fry, and 13 min for arcane. Running times for different numbers of threads are shown in Appendix C and its Fig. A3. Running times split by subcommands (e.g., barcode correction, mapping, and UMI resolution for arcane) are discussed in Appendix G.

CellRanger requires between 15.6 GB and 19.3 GB on all datasets. For the alignment-free tools, the memory usage is dominated by the *k*-mer index size. Hence, arcane, Alevin-fry and Kallisto|bustools require more memory on the three human datasets compared to the mouse dataset. Alevin-fry is by far the most memory-frugal method because of its compact minimizer-based super-*k*-mer index. It needs under 4 GB on all datasets, but it creates very large files on disk (not shown in figure). On the evaluated datasets, Kallisto|bustools requires 15.9 GB for the human dataset and 9.4 GB on the mouse dataset, thus needing less (but comparable) memory compared to CellRanger. Currently, arcane has the highest memory requirements, up to 34.7 GB on the human datasets and 26.7 GB for the mouse dataset, mainly due to the larger index size (22 GB for human and 15 GB for mouse) and because it does its computations mostly in main memory without using large temporary files.

### 4.2 arcane obtains similar quantification results as other methods

We first compared the total number of counts per cell (barcode) across tools on all four datasets (see Fig. 5 for one of the datasets and Appendix D with Figs. A4–A7 for all datasets). The total count is influenced by all steps of the quantification process: Barcode correction may increase the available number of reads per cell, the read mapping strategy may increase or decrease the number of usable reads per cell, and the UMI resolution strategy may also work both ways. arcane shows high agreement with all other approaches on all datasets. Overall, arcane shows slightly lower total counts, especially for cells with overall high counts (≥ 30 000). Alevin-fry has the highest counts compared to the other approaches, both using the `cr-like` and `parsimony-em` UMI resolution strategy due to an overall higher number of mapped reads. For example, on the `5K_Human_PBMC_v3` dataset, Alevin-fry mapped 342 831 582 reads confidently to the transcriptome, compared to 307 558 438 by Kallisto|bustools, and 307 153 285 by CellRanger. arcane mapped 285 575 960 reads, but is the only method that already discards reads from invalid barcodes before mapping.

**Fig. 5:**
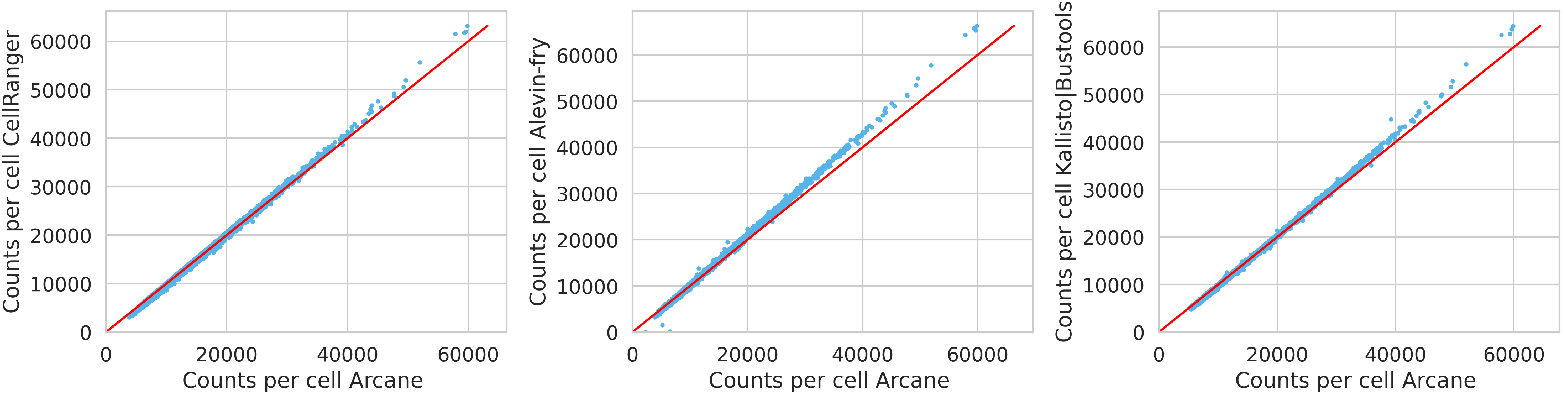
Comparison of total counts per cell (barcode) in the `5K_Human_PMBC_v3` dataset across tools (Alevin-fry in `cr-like` mode). Each blue dot corresponds to a cell. Red main diagonal: equal counts.

In more detail, we compared the number of shared cells (barcodes) and the counts in the resulting cells × genes output matrix. Table 1 presents the number of valid shared barcodes, as well as the Pearson correlation between the gene counts between approaches averaged over all shared barcodes. CellRanger has the highest number of cells (valid barcodes), followed by Alevin-fry. Both arcane and Kallisto|bustools (with UMI-tools) have a lower number of valid barcodes, which are almost a true subset of the valid barcodes of CellRanger and Alevin-fry. Among the shared barcodes, all approaches show a high Pearson correlation between their gene counts, as highlighted for arcane for the `10K_Mouse_Brain_v4` dataset in Fig. 6. Figures for the other datasets can be found in Appendix D. The `10K_Human_Melanoma_v4` is the dataset with the most variation between approaches. Alevin-fry (`cr-like`) and arcane show high correlation, but lower correlation with Kallisto|bustools and Cell-Ranger, which show high correlation with each other. In particular, in the melanoma dataset, arcane and Alevin-fry assign high counts to several genes that receive a count of zero from CellRanger and Kallisto|bustools (Appendix D, Fig. A6). This phenomenon also exists to a lesser degree in the other datasets.

**Table 1:**
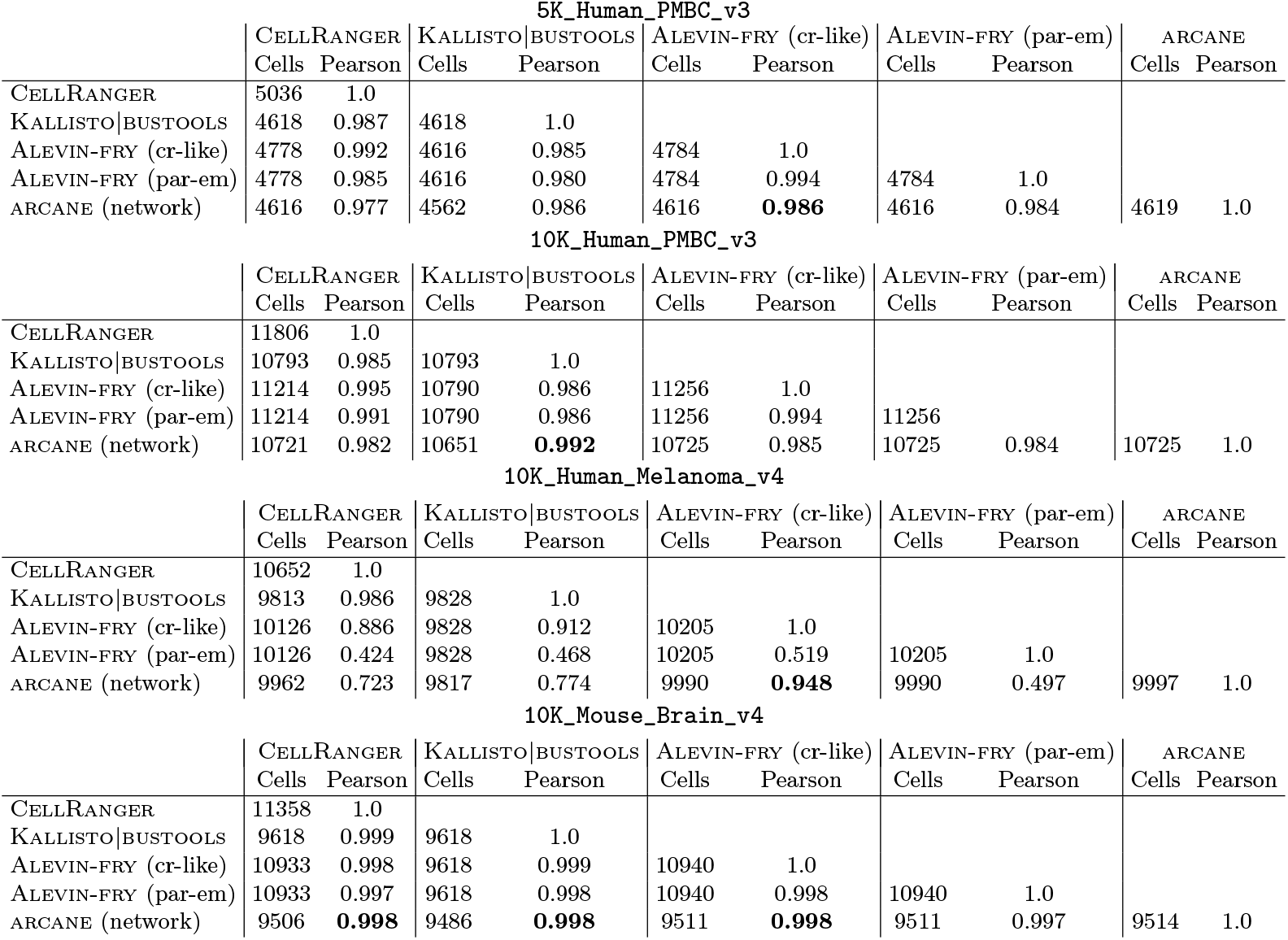
Number of shared barcodes and Pearson correlation between gene counts averaged over all shared barcodes. The bold entry shows the highest Pearson correlation of arcane with another approach.

| 5K_Human_PMBC_v3 |  |  |  |  |  |  |  |  |  |  |
| --- | --- | --- | --- | --- | --- | --- | --- | --- | --- | --- |
|  | CELLRANGER |  | KALLISTO\|BUSTOOLS |  | ALEVIN-FRY (cr-like) |  | ALEVIN-FRY (par-em) |  | ARCANE |  |
|  | Cells | Pearson | Cells | Pearson | Cells | Pearson | Cells | Pearson | Cells | Pearson |
| CELLRANGER | 5036 | 1.0 |  |  |  |  |  |  |  |  |
| KALLISTO\|BUSTOOLS | 4618 | 0.987 | 4618 | 1.0 |  |  |  |  |  |  |
| ALEVIN-FRY (cr-like) | 4778 | 0.992 | 4616 | 0.985 | 4784 | 1.0 |  |  |  |  |
| ALEVIN-FRY (par-em) | 4778 | 0.985 | 4616 | 0.980 | 4784 | 0.994 | 4784 | 1.0 |  |  |
| ARCANE (network) | 4616 | 0.977 | 4562 | 0.986 | 4616 | **0.986** | 4616 | 0.984 | 4619 | 1.0 |
| 10K_Human_PMBC_v3 |  |  |  |  |  |  |  |  |  |  |
|  | CELLRANGER |  | KALLISTO\|BUSTOOLS |  | ALEVIN-FRY (cr-like) |  | ALEVIN-FRY (par-em) |  | ARCANE |  |
|  | Cells | Pearson | Cells | Pearson | Cells | Pearson | Cells | Pearson | Cells | Pearson |
| CELLRANGER | 11806 | 1.0 |  |  |  |  |  |  |  |  |
| KALLISTO\|BUSTOOLS | 10793 | 0.985 | 10793 | 1.0 |  |  |  |  |  |  |
| ALEVIN-FRY (cr-like) | 11214 | 0.995 | 10790 | 0.986 | 11256 | 1.0 |  |  |  |  |
| ALEVIN-FRY (par-em) | 11214 | 0.991 | 10790 | 0.986 | 11256 | 0.994 | 11256 |  |  |  |
| ARCANE (network) | 10721 | 0.982 | 10651 | **0.992** | 10725 | 0.985 | 10725 | 0.984 | 10725 | 1.0 |
| 10K_Human_Melanoma_v4 |  |  |  |  |  |  |  |  |  |  |
|  | CELLRANGER |  | KALLISTO\|BUSTOOLS |  | ALEVIN-FRY (cr-like) |  | ALEVIN-FRY (par-em) |  | ARCANE |  |
|  | Cells | Pearson | Cells | Pearson | Cells | Pearson | Cells | Pearson | Cells | Pearson |
| CELLRANGER | 10652 | 1.0 |  |  |  |  |  |  |  |  |
| KALLISTO\|BUSTOOLS | 9813 | 0.986 | 9828 | 1.0 |  |  |  |  |  |  |
| ALEVIN-FRY (cr-like) | 10126 | 0.886 | 9828 | 0.912 | 10205 | 1.0 |  |  |  |  |
| ALEVIN-FRY (par-em) | 10126 | 0.424 | 9828 | 0.468 | 10205 | 0.519 | 10205 | 1.0 |  |  |
| ARCANE (network) | 9962 | 0.723 | 9817 | 0.774 | 9990 | **0.948** | 9990 | 0.497 | 9997 | 1.0 |
| 10K_Mouse_Brain_v4 |  |  |  |  |  |  |  |  |  |  |
|  | CELLRANGER |  | KALLISTO\|BUSTOOLS |  | ALEVIN-FRY (cr-like) |  | ALEVIN-FRY (par-em) |  | ARCANE |  |
|  | Cells | Pearson | Cells | Pearson | Cells | Pearson | Cells | Pearson | Cells | Pearson |
| CELLRANGER | 11358 | 1.0 |  |  |  |  |  |  |  |  |
| KALLISTO\|BUSTOOLS | 9618 | 0.999 | 9618 | 1.0 |  |  |  |  |  |  |
| ALEVIN-FRY (cr-like) | 10933 | 0.998 | 9618 | 0.999 | 10940 | 1.0 |  |  |  |  |
| ALEVIN-FRY (par-em) | 10933 | 0.997 | 9618 | 0.998 | 10940 | 0.998 | 10940 | 1.0 |  |  |
| ARCANE (network) | 9506 | **0.998** | 9486 | **0.998** | 9511 | **0.998** | 9511 | 0.997 | 9514 | 1.0 |

**Fig. 6:**
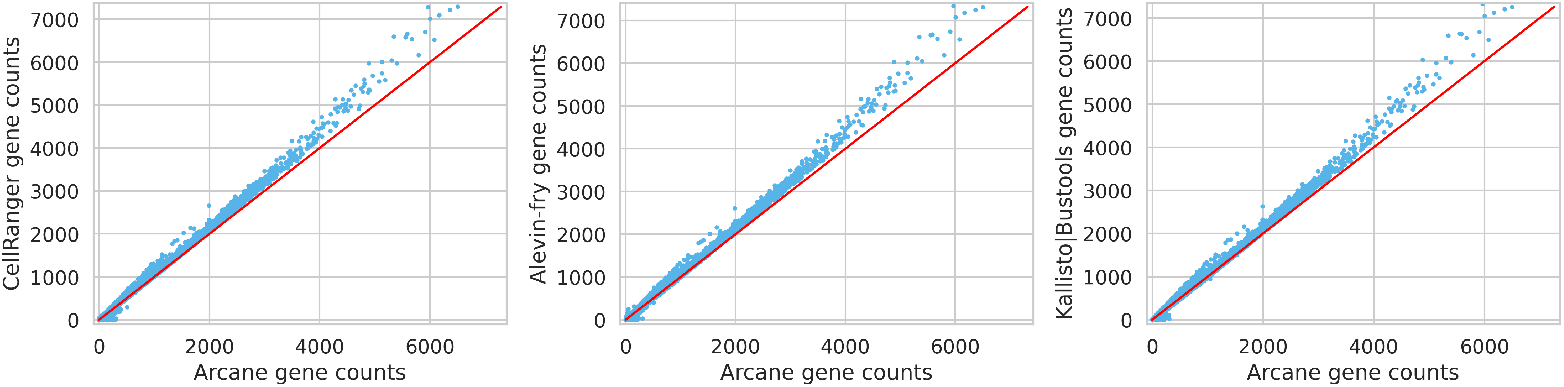
Comparison of gene quantification results between arcane and existing approaches on the `10K_Mouse_Brain_v4` dataset (Alevin-fry in `cr-like` mode). Each dot corresponds to a single gene in a single cell. The red main diagonal corresponds to equal counts.

Overall, arcane produces very similar gene quantification results compared to existing approaches, and we may assume that arcane achieves a similar accuracy and quality for downstream analysis.

## 5 Conclusion and Discussion

We presented arcane, the currently fastest method to quantify gene abundances in scRNA-seq and snRNA-seq experiments, using efficient algorithms and data structures for barcode correction and UMI resolution. We showed that storing up to three genes per gapped *k*-mer suffices to cover all genes almost completely with *k*-mers, an interesting observation by itself. Since we store the gapped *k*-mers together with their gene identifiers, we save an indirection during memory lookup, which makes arcane faster than other methods, at the cost of higher memory usage. However, arcane supports loading the index into shared memory, so several independent arcane instances may run in parallel without increasing memory usage. arcane achieves similar quantification results in comparison to existing methods with high Pearson correlation coefficients on the resulting counts in the cells × genes output matrix. Nonetheless, some genes on some datasets, especially in the human melanoma dataset, are assigned different counts by different methods, even by the two UMI resolution strategies of Alevin-fry. These differences warrant further investigation.

Further improvements to arcane may consist of additional index optimizations, increasing speed even further, but first and foremost reducing the memory requirements without sacrificing speed. Additional alternative barcode correction and UMI resolution strategies can be developed and easily integrated into the modular architecture of arcane. While the described method is applicable to all scRNA-seq or snRNA-seq data that makes use of barcode, UMI and sequence information, the current implementation is tailored to the file structure of 10X Genomics data; in the future, we will also support other formats. Currently, arcane uses 16-bit gene IDs which allow us to encode up to 65 536 different genes, including protein-coding and lncRNA genes. If a more fine-grained classification is desired, the gene bits can easily be increased, resulting in a higher memory requirement. In addition, arcane is currently limited to outputting merged gene counts. Thus, future work will entail the option to separate counts into spliced and unspliced, making arcane also applicable to RNA velocity analysis.

## Appendix

## Appendix A: Knee method after barcode correction

Figure A1 shows so-called knee plots with complementary cumulative distribution functions, with flipped log-scaled axes: For a given read count (on the log-scaled y-axis), the cumulative number of barcodes (on the log-scaled x-axis) that have at least the given count is shown. Since we apply this method after barcode correction but before UMI resolution, we consider the total number of reads for each barcode, in contrast to other methods that consider the number of distinct UMIs. The vertical red line shows where UMI-tools places the knee (used in arcane). In subsequent steps, arcane only considers the frequent barcodes to the left of the knee to be valid cells.

**Fig. A1:**
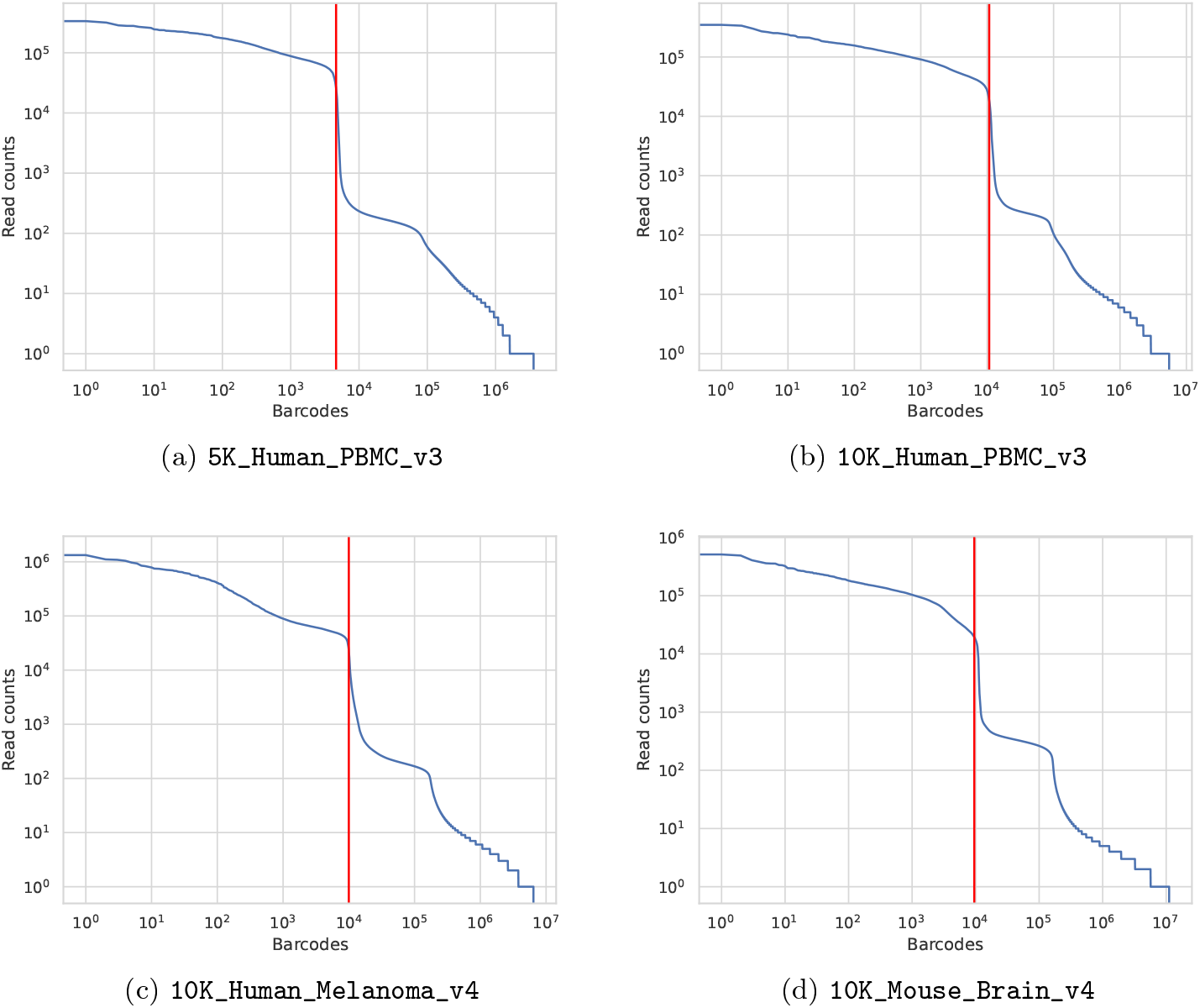
Knee plots showing the number of cells (barcodes, on the x-axis, log-scaled) that have at least the given read count (y-axis, log-scaled) after barcode correction. The vertical red line shows the threshold below which arcane discard barcodes due to a too low read count. Each of the four panels shows one of the investigated datasets.

**Fig. A2:**
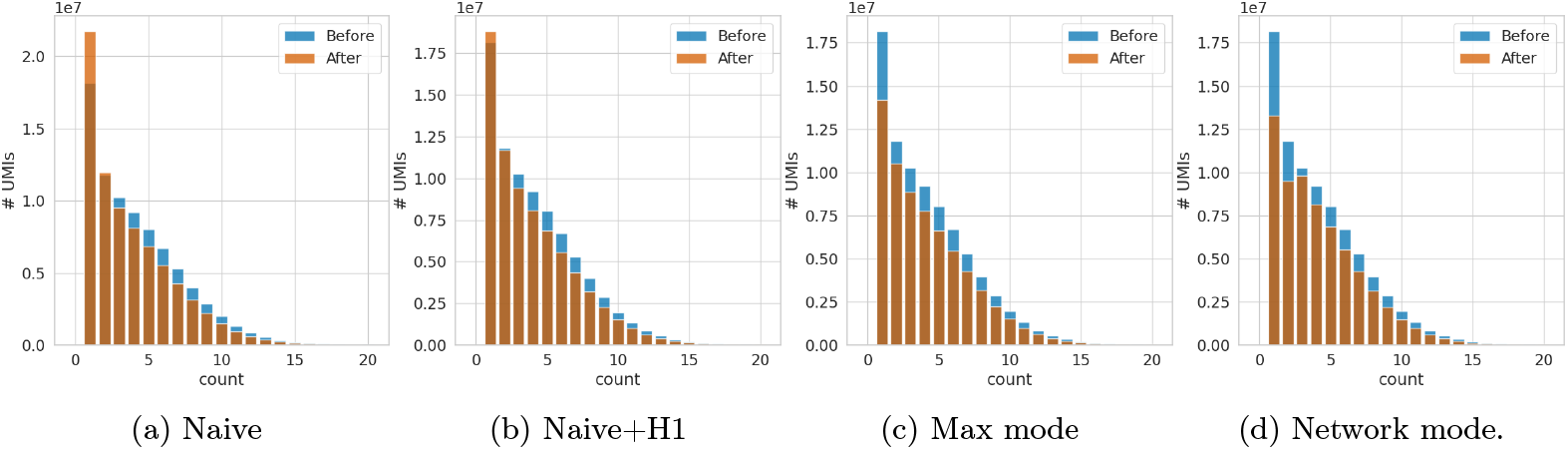
UMI counts before (blue) and after (orange) UMI resolution using different resolution strategies on the `5K_PBMC_V3` data set.

## Appendix B: UMI resolution

Figure A2 shows the effect of different UMI resolution strategies on the UMI histogram on the `5K_PBMC_V3` dataset. In all four histograms, the blue histogram (before resolution), shows the raw UMI count histogram, accumulated over all cells (retained valid barcodes). The orange histograms show the UMI counts after resolution, i.e, the UMI count for each incremented gene.

From the shown `5K_PBMC_V3` dataset, using retained valid barcodes, 285 575 960 reads were uniquely mapped and 63 138 181 were unmapped, either due to multi-mapping or because no gapped *k*-mer of the read was found in the index. While the blue histograms (before) include UMIs for unmapped reads, the orange histograms (after) do not.

The UMI resolution strategies described below exist in addition to the Network mode described in the main article. Both the Max mode and the Network mode (Sec. 3.5 in the main article) result in similar histograms on the `5K_PBMC_V3` dataset and reduce the number of singleton UMIs (count of 1). The resulting count is closer to the expected proportion under a Poisson model.

### Naive mode

In naive mode, UMIs (with the same barcode) are only collapsed if they share the exact same barcode and are mapped to the same gene. Thus, a UMI with a high count *c* from two or more genes will be split into several (UMI, gene) pairs with smaller counts that sum to *c*. Therefore, the number of singletons increases after naive UMI resolution.

### Naive+H1 mode

UMIs are collapsed if they map to the same barcode, are in the same connected component in the graph with edges between Hamming distance 1 neighbors and map to the same gene. However, this approach still does not solve the problem of many singletons. This may be caused by wrong gene identifications for low quality reads or UMIs with a Hamming distance *>* 1 to the correct UMI.

### Max mode

Given a connected component in the Hamming-distance-1 neighbor graph, we collect the genes associated with any of the UMIs in the component and, for each gene, add the counts across all UMIs in the component. Then we assume that the component only represents the gene *g*^∗^ with the highest summed UMI count (similar approach to CellRanger). This mode is robust against erroneous UMIs but may underestimate the total number of unique UMIs if two true UMIs in the sample have a Hamming distance of 1 or if the same UMI is (randomly) used twice for different molecules.

## Appendix C: Running times for different numbers of threads

Figure A3 shows the wall clock running time for CellRanger, Alevin-fry, Kallisto|bustools and arcane on the `10K_Human_PBMC_v3` datasets for different number of threads, showing that the wall time decreases for increasing number of threads for all methods. For all number of threads, arcane is the fastest method.

## Appendix D: Correlation between tools

Figures A4–A7, one figure per dataset, show the degree of agreement between different tools, considering the total gene counts in a cell and gene counts for a single gene in a single cell. In each figure, the top part (A) shows total counts per cell: Each dot represents a barcode (cell), and x-axis and y-axis show counts from different tools. All approaches yield similar total gene counts, with points mostly clustering around the main diagonal, with other tools showing a slightly steeper slope than arcane. The bottom part (B) shows counts per gene per cell: Each dot represents a barcode-gene combination (*b, g*) and shows the resulting counts *X*_*b,g*_ for cell (barcode) *b* and gene *g* across different approaches. While the diagonal is clearly visible, there are count values that differ significantly between approaches. In particular, the human melanoma dataset shows many differences between all methods, which require further investigation.

**Fig. A3:**
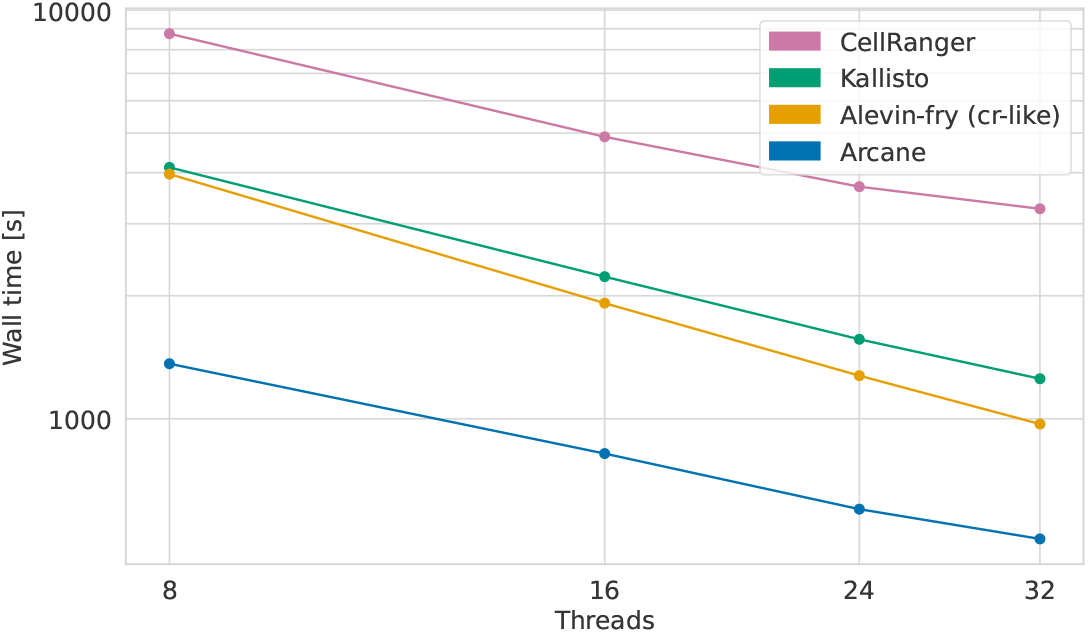
Wall clock running times (log scale) in seconds measured for 8, 16, 24 and 32 threads (log-scale) on the `10K_Human_PBMC_v3` dataset.

## Appendix E: Code and Data Availability

Source code of arcane is available online at https://gitlab.com/rahmannlab/ARCANE. All used datasets were produced by 10X Genomics and are available from their website:

~~~
5K_Human_PBMC_v3 https://support.10xgenomics.com/single-cell-gene-expression/datasets/3.0.2/5k_pbmc_v3
10K_Human_PBMC_v3 https://support.10xgenomics.com/single-cell-gene-expression/datasets/3.0.0/pbmc_10k_v3
10K_Human_Melanoma_v4 https://www.10xgenomics.com/datasets/10k-human-dtc-melanoma-GEM-X
10K_Mouse_Brain_v4 https://www.10xgenomics.com/datasets/10k-Mouse-Brain-CNIK-3p-gemx
~~~

## Appendix F: Implementation details

arcane is implemented as a workflow-friendly command line tool in Python, using `numba` to just-in-time compile the code.

### Parallelization

We use a consumer-producer model to parallelize the mapping step across many threads. The threads are organized into three levels and communicate via shared buffers. The threads in the first level read the (*b, m, s*) triples that are in practice separated into two files and write the sequence information into a buffer. The threads in the second level consume the (*b, m, s*) buffers and perform the mapping of *s* to a gene identifier *g* using *k*-mer lookups. The resulting (*b, m, g*) triples are written into output buffers that are consumed by the thread of the third level, which inserts the (*m, g*) pairs into the *b*-block of the UMI-gene array. Since we can encode the UMI *m* (assuming a length of |*m*| = 12) with 24 bits and the gene ids with 16 bits, we can store the (*m, g*) pair in a single 64-bit integer.

### Index in shared memory

For multi-sample scenarios, arcane supports loading the index into shared memory. Since the mapping only needs read-only access to the index, multiple independent arcane instances can access the same index to map the reads. This has the advantage that the memory footprint does not increase if multiple arcane instances are run in parallel.

**Fig. A4:**
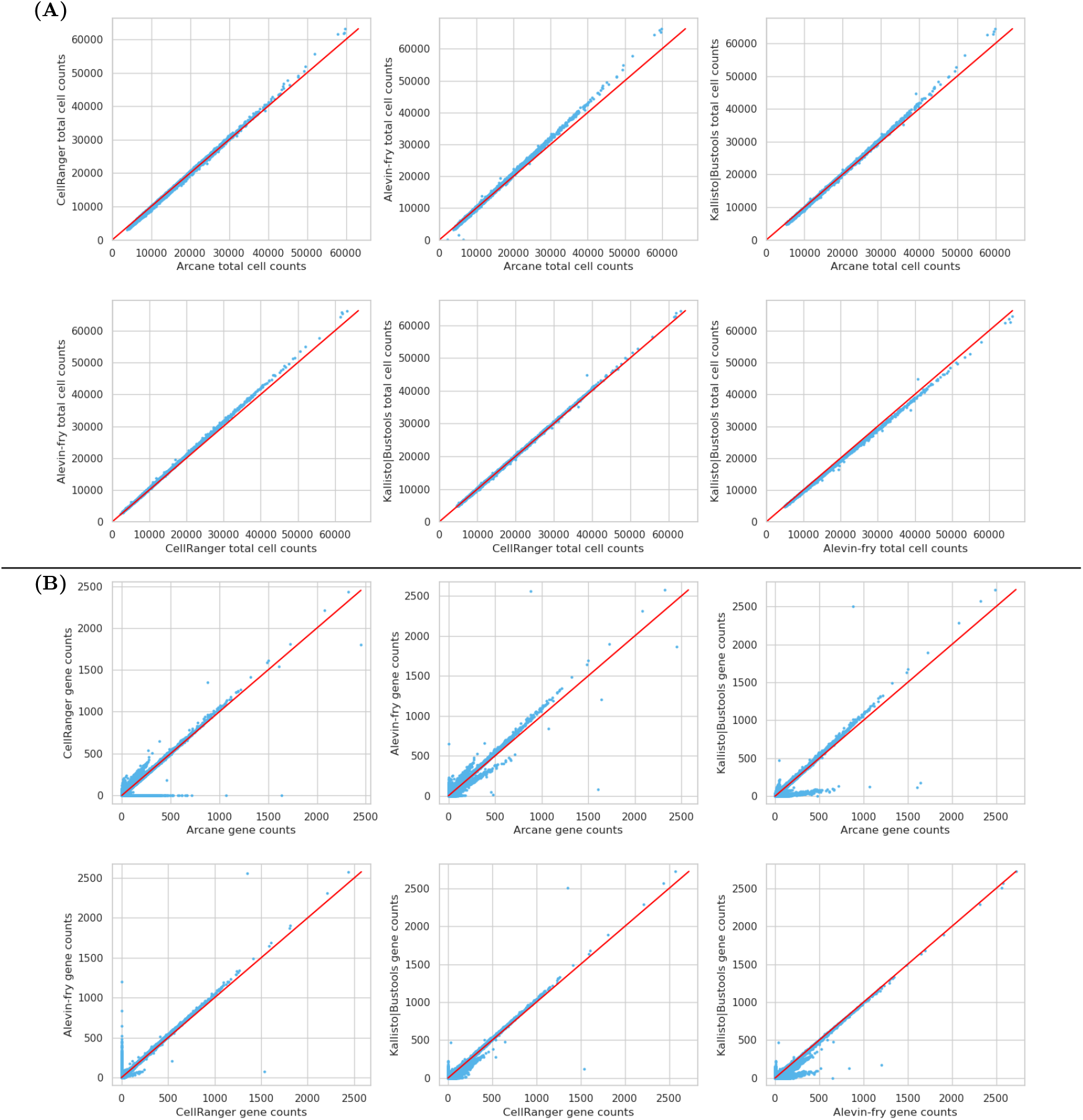
Pairwise correlation between different approaches on the `5K_Human_PBMC_v3` dataset. **A** Each dot corresponds to a single barcode. The (*x, y*) coordinates correspond to the total gene counts for this barcode in both methods (for the barcode all gene abundances are summed up). **B** Each dot corresponds to a single gene in a single barcode. The (*x, y*) coordinates correspond to the gene count in this barcode for both methods.

**Fig. A5:**
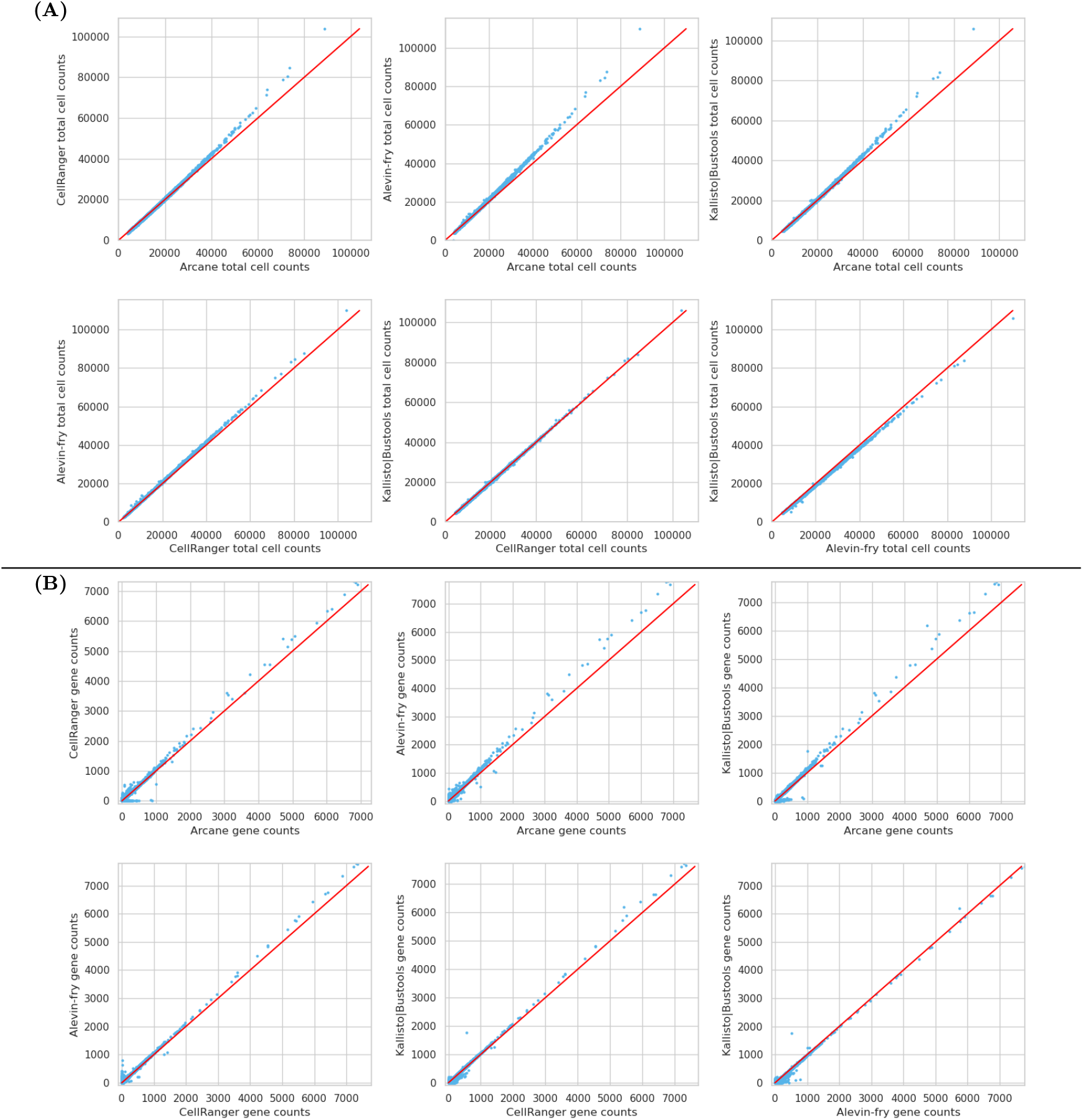
Pairwise correlation between different approaches on the `10K_Human_PBMC_v3` dataset. **A** Each dot corresponds to a single barcode. The (*x, y*) coordinates correspond to the total gene counts for this barcode in both methods (for the barcode all gene abundances are summed up). **B** Each dot corresponds to a single gene in a single barcode. The (*x, y*) coordinates correspond to the gene count in this barcode for both methods.

**Fig. A6:**
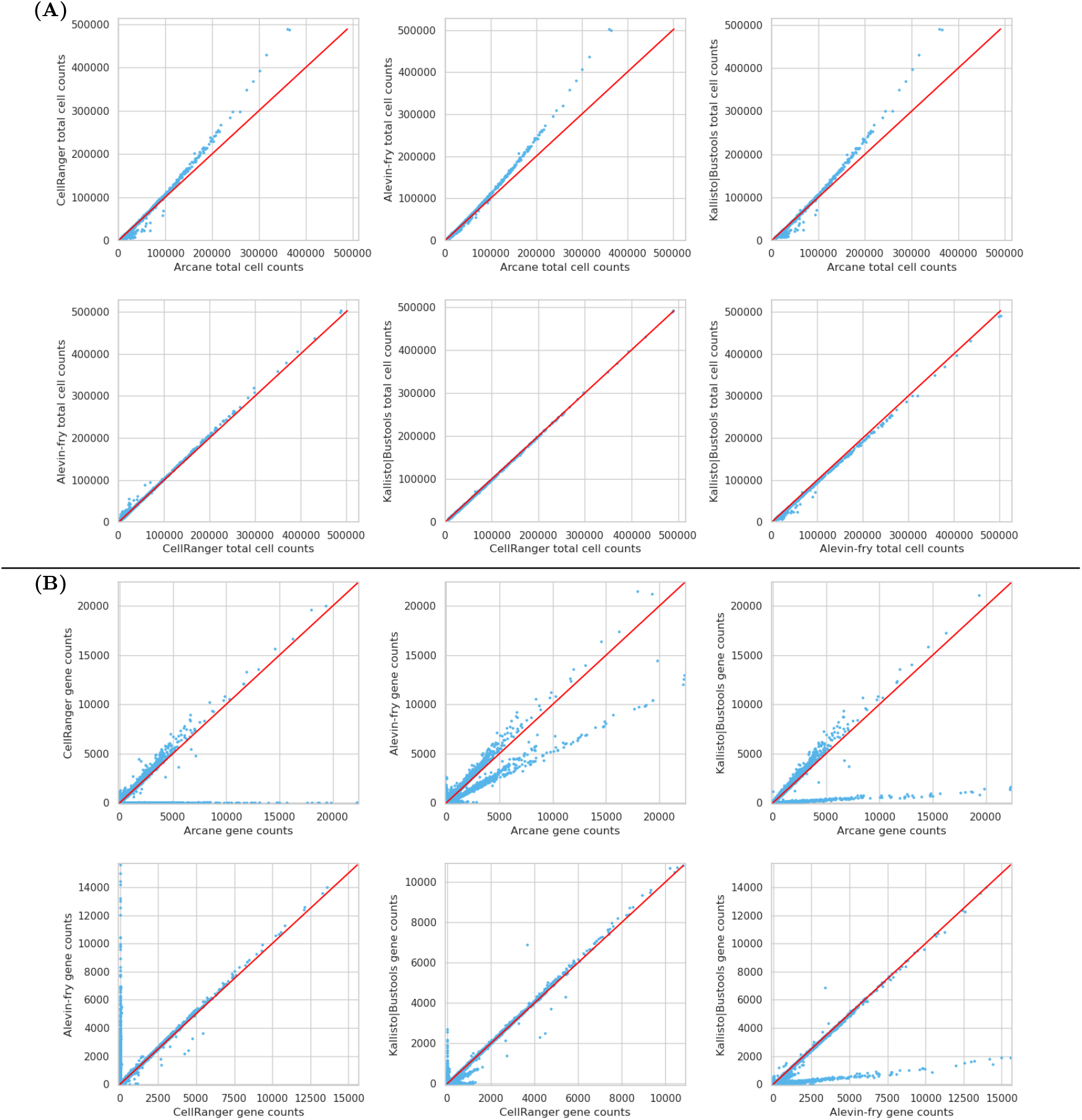
Pairwise correlation between different approaches on the `10K_Human_Melanoma_v4` dataset. **(A)** Each dot corresponds to a single barcode. The (*x, y*) coordinates correspond to the total gene counts for this barcode in both methods (for the barcode all gene abundances are summed up). **(B)** Each dot corresponds to a single gene in a single barcode. The (*x, y*) coordinates correspond to the gene count in this barcode for both methods.

**Fig. A7:**
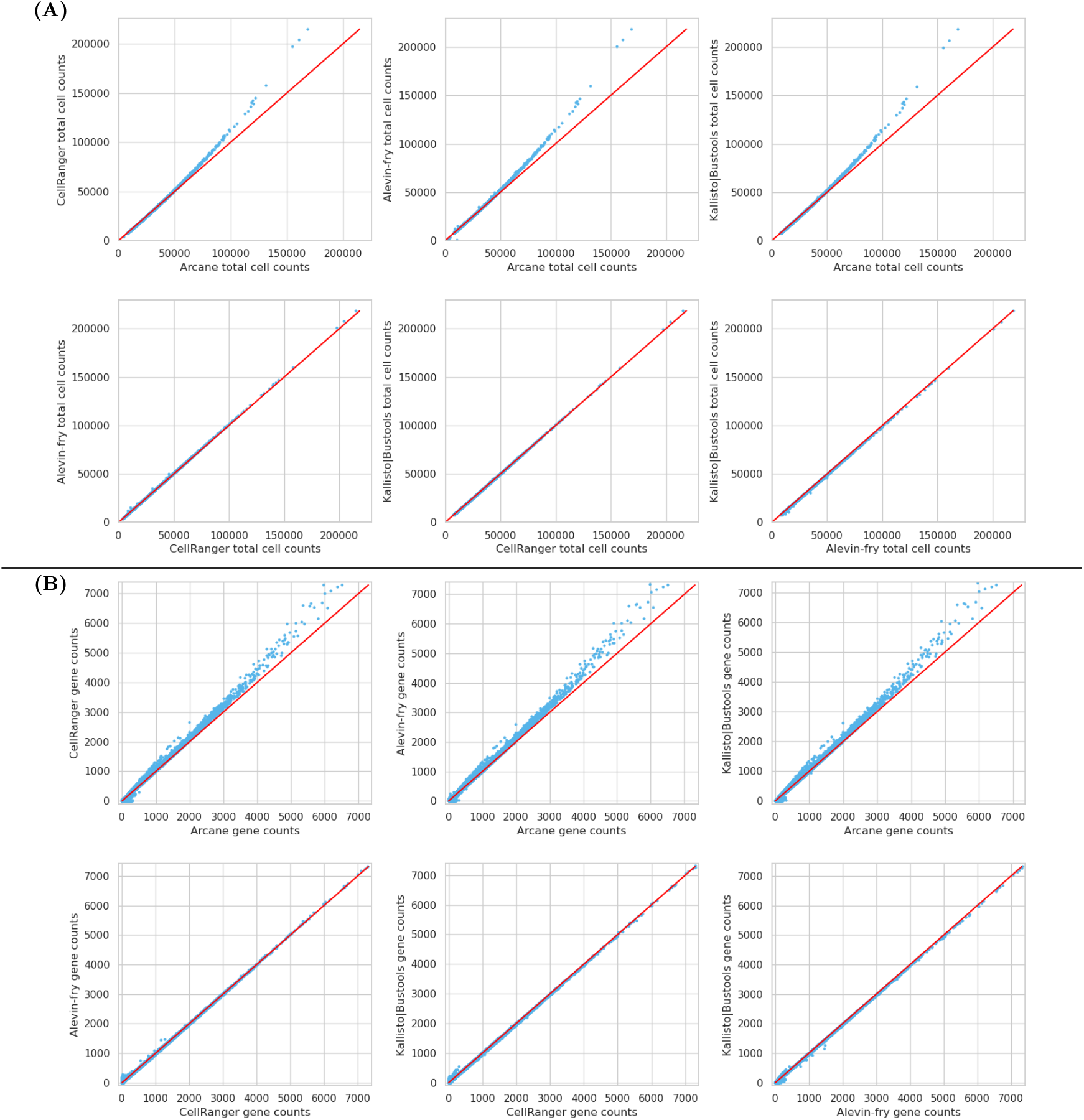
Pairwise correlation between different approaches on the `10K_Mouse_Brain_v4` dataset. **(A)** Each dot corresponds to a single barcode. The (*x, y*) coordinates correspond to the total gene counts for this barcode in both methods (for the barcode all gene abundances are summed up). **(B)** Each dot corresponds to a single gene in a single barcode. The (*x, y*) coordinates correspond to the gene count in this barcode for both methods.

## Appendix G: Running time for each step

### Appendix G:.1 arcane

arcane consists of three main steps: (1) barcode correction, (2) mapping, and (3) UMI resolution. On the `5K_Human_PBMC_v3` dataset (using 16 threads), the barcode correction takes 1 min 25 s, the mapping 5 min 29 s and the UMI resolution 24 s. The total running time is thus 7 min 18 s.

### Appendix G:.2 Alevin-fry

To quantify gene expression with Alevin-fry, four Alevin-fry subcommands are executed: (1) mapping, (2) generating a permit (positive) list, (3) collation, and (4) quantification. The mapping takes 16 min 38 s, generating the permit list takes 14 s, and both collation and quantification take 8 s for the `5K_Human_PBMC_v3` dataset using 16 threads, resulting in a total running time of 16 min 53 s. Note that the mapping step also produces the intermediate RAD files, which allows faster processing in the subsequent steps.

### Appendix G:.3 Kallisto|bustools

Kallisto|bustools internally performs six steps, listed below together with the running time on the `5K_Human_PBMC_v3` dataset (using 16 threads).

1. `kallisto bus`: 20 min 46 s
2. `bustools sort`: 42 s
3. `bustools inspect`: 15 s
4. `bustools correct`: 16 s
5. `bustools sort`: 19 s
6. `bustools count`: 18 s

The total running time is 22 min 36 s.

### Appendix G:.4 CellRanger

CellRanger does not output information about the running time of different steps. The total running time on the `5K_Human_PBMC_v3` dataset using 16 threads is 50 min 17 s.

### Appendix G:.5 Summary

Although there is no one-to-one correspondence between tools, we see that the mapping step of arcane is much faster compared to the other methods. The barcode correction and quantification of Alevin-fry are faster than the steps in arcane, partly because generating the intermediate RAD format enables faster processing later on.

## Footnotes

1 Even for an error rate of 10% per bp for a barcode of length *n* = 16, this is true: From the binomial distribution with *n* = 16 and *p* = 0.1, we see that 18.5% of barcodes are error-free, while 32.9% have one error, so each of these 16 distinct molecules has a frequency of 2.06% and is 9 times less frequent than the correct barcode (assuming randomness of errors).

2 The term *color* is common in the literature; it abstracts away from the concrete type of objects stored; in this case, we store numerical gene identifiers, so a color represents a gene, and a color set represents a set of genes.

3 If barcodes are much longer, different techniques than bit arrays must be considered.

## Notes

https://gitlab.com/rahmannlab/arcane

### Competing Interest Statement

The authors have declared no competing interest.

### Summary of Updates

Small improvements and extended time evaluation.

## References

1. Almodaresi, F., Sarkar, H., Srivastava, A., Patro, R.: A space and time-efficient index for the compacted colored de Bruijn graph. Bioinformatics 34(13), i169–i177 (2018). 10.1093/bioinformatics/bty292

2. Bray, N.L., Pimentel, H., Melsted, P., Pachter, L.: Near-optimal probabilistic RNA-seq quantification. Nature Biotechnology 34(5), 525–527 (2016). 10.1038/nbt.3519

3. Brinda, K., Sykulski, M., Kucherov, G.: Spaced seeds improve k-mer-based metagenomic classification. Bioinformatics 31(22), 3584–3592 (2015). 10.1093/bioinformatics/btv419

4. Compeau, P.E.C., Pevzner, P.A., Tesler, G.: How to apply de Bruijn graphs to genome assembly. Nature Biotechnology 29(11), 987–991 (2011). 10.1038/nbt.2023

5. Davis, R.T., Blake, K., Ma, D., Gabra, M.B.I., Hernandez, G.A., Phung, A.T., Yang, Y., Maurer, D., Lefebvre, A.E.Y.T., Alshetaiwi, H., Xiao, Z., Liu, J., Locasale, J.W., Digman, M.A., Mjolsness, E., Kong, M., Werb, Z., Lawson, D.A.: Transcriptional diversity and bioenergetic shift in human breast cancer metastasis revealed by single-cell RNA sequencing. Nature Cell Biology 22(3), 310–320 (2020). 10.1038/s41556-020-0477-0

6. Dobin, A., Davis, C.A., Schlesinger, F., Drenkow, J., Zaleski, C., Jha, S., Batut, P., Chaisson, M., Gingeras, T.R.: STAR: ultrafast universal RNA-seq aligner. Bioinformatics 29(1), 15–21 (2013). 10.1093/bioinformatics/bts635

7. Fleming, S.J., Chaffin, M.D., Arduini, A., Akkad, A.D., Banks, E., Marioni, J.C., Philippakis, A.A., Ellinor, P.T., Babadi, M.: Unsupervised removal of systematic background noise from droplet-based single-cell experiments using CellBender. Nature Methods 20(9), 1323–1335 (2023). 10.1038/s41592-023-01943-7

8. González, R., Grabowski, S., Mäkinen, V., Navarro, G.: Practical implementation of rank and select queries. In: Poster Proc. Volume of 4th Workshop on Efficient and Experimental Algorithms (WEA). pp. 27–38. CTI Press and Ellinika Grammata Greece (2005)

9. He, D., Patro, R.: simpleaf: a simple, flexible, and scalable framework for single-cell data processing using alevin-fry. Bioinformatics 39(10), btad614 (2023). 10.1093/bioinformatics/btad614

10. He, D., Zakeri, M., Sarkar, H., Soneson, C., Srivastava, A., Patro, R.: Alevin-fry unlocks rapid, accurate and memory-frugal quantification of single-cell RNA-seq data. Nature Methods 19(3), 316–322 (2022). 10.1038/s41592-022-01408-3

11. Huang, D., Ma, N., Li, X., Gou, Y., Duan, Y., Liu, B., Xia, J., Zhao, X., Wang, X., Li, Q., Rao, J., Zhang, X.: Advances in single-cell RNA sequencing and its applications in cancer research. Journal of Hematology & Oncology 16(1), 98 (2023). 10.1186/s13045-023-01494-6

12. Iqbal, Z., Caccamo, M., Turner, I., Flicek, P., McVean, G.: De novo assembly and genotyping of variants using colored de Bruijn graphs. Nature Genetics 44(2), 226–232 (2012). 10.1038/ng.1028

13. Li, X., Wang, C.Y.: From bulk, single-cell to spatial RNA sequencing. International Journal of Oral Science 13(1), 36 (2021). 10.1038/s41368-021-00146-0

14. Ma, B., Tromp, J., Li, M.: PatternHunter: faster and more sensitive homology search. Bioinformatics 18(3), 440–445 (2002). 10.1093/bioinformatics/18.3.440

15. Macosko, E.Z., Basu, A., Satija, R., Nemesh, J., Shekhar, K., Goldman, M., Tirosh, I., Bialas, A.R., Kamitaki, N., Martersteck, E.M., Trombetta, J.J., Weitz, D.A., Sanes, J.R., Shalek, A.K., Regev, A., McCarroll, S.A.: Highly Parallel Genome-wide Expression Profiling of Individual Cells Using Nanoliter Droplets. Cell 161(5), 1202–1214 (2015). 10.1016/j.cell.2015.05.002

16. Melsted, P., Booeshaghi, A.S., Liu, L., Gao, F., Lu, L., Min, K.H.J., da Veiga Beltrame, E., Hjörleifsson, K.E., Gehring, J., Pachter, L.: Modular, efficient and constant-memory single-cell RNA-seq preprocessing. Nature Biotechnology 39(7), 813–818 (2021). 10.1038/s41587-021-00870-2

17. Pibiri, G.E.: Sparse and skew hashing of k-mers. Bioinformatics 38, i185–i194 (2022). 10.1093/bioinformatics/btac245

18. Rood, J.E., Maartens, A., Hupalowska, A., Teichmann, S.A., Regev, A.: Impact of the Human Cell Atlas on medicine. Nature Medicine 28(12), 2486–2496 (2022). 10.1038/s41591-022-02104-7

19. Shainer, I., Stemmer, M.: Choice of pre-processing pipeline influences clustering quality of scRNA-seq datasets. BMC Genomics 22(1), 661 (2021). 10.1186/s12864-021-07930-6

20. Smith, T., Heger, A., Sudbery, I.: UMI-tools: modeling sequencing errors in unique molecular identifiers to improve quantification accuracy. Genome Research 27(3), 491–499 (2017). 10.1101/gr.209601.116

21. Srivastava, A., Malik, L., Smith, T., Sudbery, I., Patro, R.: Alevin efficiently estimates accurate gene abundances from dscRNA-seq data. Genome Biology 20(1), 65 (2019). 10.1186/s13059-019-1670-y

22. Sullivan, D.K., Min, K.H.J., Hjörleifsson, K.E., Luebbert, L., Holley, G., Moses, L., Gustafsson, J., Bray, N.L., Pimentel, H., Booeshaghi, A.S., Melsted, P., Pachter, L.: kallisto, bustools and kb-python for quantifying bulk, single-cell and single-nucleus RNA-seq. Nature Protocols 20(3), 587–607 (2025). 10.1038/s41596-024-01057-0

23. Tarjan, R.E.: A class of algorithms which require nonlinear time to maintain disjoint sets. Journal of Computer and System Sciences 18(2), 110–127 (1979). 10.1016/0022-0000(79)90042-4

24. Tirosh, I., Suva, M.L.: Cancer cell states: Lessons from ten years of single-cell RNA-sequencing of human tumors. Cancer Cell 42(9), 1497–1506 (2024-09-09). 10.1016/j.ccell.2024.08.005

25. Zentgraf, J., Rahmann, S.: Fast lightweight accurate xenograft sorting. Algorithms for Molecular Biology 16(1), 2 (2021). 10.1186/s13015-021-00181-w

26. Zentgraf, J., Rahmann, S.: Fast Gapped k-mer Counting with Subdivided Multi-Way Bucketed Cuckoo Hash Tables. In: 22nd International Workshop on Algorithms in Bioinformatics (WABI 2022). vol. 242, pp. 12:1–12:20 (2022). 10.4230/LIPIcs.WABI.2022.12

27. Zentgraf, J., Rahmann, S.: Swiftly identifying strongly unique k-mers. In: Pissis, S.P., Sung, W. (eds.) 24th International Workshop on Algorithms in Bioinformatics, WABI 2024, September 2-4, 2024, Royal Holloway, London, United Kingdom. LIPIcs, vol. 312, pp. 15:1–15:15. Schloss Dagstuhl - Leibniz-Zentrum für Informatik, Germany (2024). 10.4230/LIPICS.WABI.2024.15

28. Zentgraf, J., Rahmann, S.: Design of Worst-Case-Optimal Spaced Seeds. In: Brejová, B., Patro, R. (eds.) 25th International Conference on Algorithms for Bioinformatics (WABI 2025). Leibniz International Proceedings in Informatics (LIPIcs), vol. 344, pp. 22:1–22:17. Schloss Dagstuhl – Leibniz-Zentrum für Informatik (2025). 10.4230/LIPIcs.WABI.2025.22

29. Zentgraf, J., Rahmann, S.: Swiftly identifying strongly unique k-mers. Algorithms for Molecular Biology 20(1), 13 (2025). 10.1186/s13015-025-00286-6

30. Zentgraf, J., Schmitz, J.E., Rahmann, S.: Cleanifier: Contamination removal from microbial sequences using spaced seeds of a human pangenome index. Bioinformatics (2025). 10.1093/bioinformatics/btaf632

31. Zheng, G.X.Y., Terry, J.M., Belgrader, P., Ryvkin, P., Bent, Z.W., Wilson, R., Ziraldo, S.B., Wheeler, T.D., McDermott, G.P., Zhu, J., Gregory, M.T., Shuga, J., Montesclaros, L., Underwood, J.G., Masquelier, D.A., Nishimura, S.Y., Schnall-Levin, M., Wyatt, P.W., Hindson, C.M., Bharadwaj, R., Wong, A., Ness, K.D., Beppu, L.W., Deeg, H.J., McFarland, C., Loeb, K.R., Valente, W.J., Ericson, N.G., Stevens, E.A., Radich, J.P., Mikkelsen, T.S., Hindson, B.J., Bielas, J.H.: Massively parallel digital transcriptional profiling of single cells. Nature Communications 8(1), 14049 (2017). 10.1038/ncomms14049

